# Revisiting the effect of population size on cumulative cultural evolution

**DOI:** 10.1101/001529

**Authors:** Ryan Baldini

## Abstract

Henrich (2004) argued that larger populations can better maintain complex technologies because they contain more highly skilled people whom others can imitate. His original model, however, did not distinguish the effects of population size from population density or network size; a learner’s social network included the entire population. Does population size remain important when populations are subdivided and networks are realistically small? I use a mathematical model to show that population size has little effect on *equilibrium* levels of mean skill under a wide range of conditions. The effects of network size and transmission error rate usually overshadow that of population size. Population size can, however, affect the *rate* at which a population approaches equilibrium, by increasing the rate at which innovations arise. This effect is small unless innovation is very rare. Whether population size predicts technological complexity in the real world, then, depends on whether technological evolution is innovation-limited and short of equilibrium. The effect of population “connectedness,” via migration or trade, is similar. I discuss the results of this analysis in light of the current empirical debate.

## 1 Introduction

Henrich (2004) used a mathematical model to argue that large populations better facilitate cumulative cultural evolution, i.e. the process by which human populations acquire more information, higher skill levels for complex tasks, and better technologies through time. Others had proposed similar ideas before (Shennan, 2001), but Henrich’s mechanistic hypothesis was perhaps more original and compelling. The argument is simple: larger populations contain more exceptionally skilled individuals whom others can copy. Above a critical population size, the positive effect of imitating the skillful becomes large enough to cancel out the negative effect of transmission error, leading to increasingly advanced skills and technologies. Henrich originally used this theory to explain the prehistoric loss of technological complexity among Tasmanian hunter-gatherers following their isolation from the Australian mainland. This idea was contentious: see Read’s (2006; 2011) critiques and Henrich’s (2006) rebuttal.

Regardless of the original empirical intent, however, the idea has since taken off. Theorists have built variations on the original model that include population structure (Powell et al., 2009, 2010), alternative social learning biases (Aoki et al., 2011; Bentley and O’brien, 2011; Vaesen, 2012), various forms of transmission error (Mesoudi, 2011; Vaesen, 2012), overlapping generations and small social networks (Kobayashi and Aoki, 2012), and trade networks in economic contexts (Kelly, 2009). Some have applied the model to particular prehistoric contexts (Powell et al., 2009, 2010; Lycett and Norton, 2010; Marquet et al., 2012). Empirical tests of the theory includes small-scale lab experiments (Caldwell and Millen, 2010; Derex et al., 2013; Muthukrishna et al., 2014) and the search for a relationship between population size and technological complexity in real human populations. Papers of the latter style claim to find both supporting (Kline and Boyd, 2010; Collard et al., 2013b) and disputing or ambivalent (Collard et al., 2005, 2013a; Read, 2008, 2012) evidence for the theory.

The purpose of this paper is to distinguish the relative effects of population size and social network size on cumulative cultural evolution. Henrich’s original model assumed that *every* learner could identify the single most skilled individual in the entire population, implying that one’s social network size (the number of people from whom one could learn) is equal to the total population size. This is clearly unrealistic for any human population. Does population size remain predictive of cultural complexity when social networks are realistically small? Furthermore, how do population size and density separately affect cultural evolution? Does simply increasing a population’s range, e.g. by adding new subpopulations to its edge, have the same effect as increasing the density of its subpopulations? This question is important because these variables may be confounded in empirical studies.

I use a mathematical model similar to Henrich’s to show that population size usually has very little effect on equilibrium levels of mean skill. Small changes in social network size, innovation rate, or transmission error rate greatly alter equilibrium skill, suggesting that cross-cultural variation in these parameters should swamp the effects of population size. Population size can, however, strongly affect the rate at which the population approaches equilibrium, because larger populations discover new innovations more often. This effect is strongest when innovation is very rare because, in that case, the total rate of innovation in the population is approximately proportional to population size. Whether population size predicts technological complexity in the reality depends on whether observed levels of technological complexity represent equilibria, or whether they are just snapshots of a slow process of growth.

I also show the relative effects of population density (as opposed to total population size) and migration between subpopulations. I find that, unlike population size, density can noticeably affect equilibrium skill levels. This effect is still relatively small compared to that of network size - unless learners can sample the entire subpopulation, in which case network size and population density are equivalent. Migration between subpopulations has little effect on equilibrium. Like population size, however, it can increase the rate of skill growth by making rare innovations available to all subpopulations. Hence population “connectedness,” like population size, is most important when innovations are rare and populations have not converged on equilibria.

The next section constructs the model. Section 3 then attempts to explain the basic results of the model mathematically. Section 4 largely confirms these explanations with simulation results. Section 5 discusses how my results may clarify disagreements in the empirical literature.

## 2 The model

A population of socially learning organisms is divided into *M* subpopulations, each of size *N*. This allows us to distinguish the effects of total population size (*N × M*) from population density (*N*). Each learning round, individuals sample *n* others from the local subpopulation to form a social network, and imitate the most skilled individual in their networks, as detailed below. A proportion *m* of the population then migrates to a different subpopulation. This model structure can apply to both short timescales, where individuals repeatedly learn and move about within their lifetime, and longer timescales, where individuals learn only once and social learning occurs across generations.

An individual’s skill level, *z*, is the proportion of correct “decisions” or techniques he employs, out of a total number of decisions *d*. One’s decision vector shows which decisions are correct and which are incorrect, denoted respectively by 1 and 0. For example, an individual with decision vector 100101… is correct with respect to decisions one, four and six, and incorrect for two, three, and five, and so on. Thus *z* ∈ {0, 1/*d*, 2/*d*, …, (*d* − 1)/*d*, 1}; *z* = 1 implies that the individual makes all correct decisions, and *z* = 0 implies all incorrect decisions. The convenience of this formulation lies in the natural maximum and minimum values for *z*; unlike Henrich’s original model, for example, indefinite skill growth or decline is impossible. Although one can imagine cases where this is a useful representation of skill in some task (e.g. a farmer’s success in crop production depends on such decisions as what, where, when, and how to plant), I use it mainly for convenience; I do not mean to imply that this model is any more realistic than others in the literature.

Learners copy the decision vector of the most skilled individual in their network. Transmission error, however, ensures that learners rarely acquire the same vector as their model. Sometimes a learner acquires the wrong decision even if their model employed the correct one, and vice versa. The deleterious error rate is denoted by *u* and the positive innovation rate by *v*. I assume *u ≥ v* throughout, so that deleterious transmission errors are more common that beneficial ones. This captures Henrich’s original assumption that transmission error tends to erode mean skill.

Like previous models, mine optimistically assumes that learners always identify and copy the most skilled individuals in their networks. This is unrealistic: identifying a person’s skill in complex tasks is difficult (Hill and Kintigh, 2009). Uncertainty in payoffs greatly reduce the effectiveness of the success-biased learning strategy modeled here (Baldini, 2013), so including uncertainty will probably lead to lower mean skill levels. I doubt, however, that the general conclusions of this paper depend on this assumption.

## 3 The effects of population size: mathematical arguments

The complexity of this model preludes much mathematical analysis. Although it resembles quantitative genetic models of directional selection under haploid inheritance and finite population size, the classic results of that literature generally fail to apply here. The reason is that selection, being driven by decision-making forces, can be very strong in this model, and the mutation rate, analogous here to transmission error and innovation, can be very large.

Diffusion methods, which provide the most tractable results for finite population sizes, require the assumption of weak evolutionary forces and are therefore inappropriate. Furthermore, the effective lack of “cultural recombination” in this model implies appreciable correlation between different decisions at equilibrium, rendering single-locus models of genetic evolution inaccurate. Thus, this section provides only the simplest, isolated mathematical arguments, which provide some grasp on the full dynamics.

### 3.1 Total population size affects the probability that innovations arise

Total population size, *NM*, most strongly affects the rate at which innovations appear in the population. This effect is largest when innovation is very rare, specifically *υ* ≪ 1*/*(*NM*). To see this, consider the probability that a novel innovation arises for some decision in a population of size *NM*. Call this probability *P*. Then *P* is

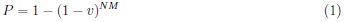

The expected number of new innovations is this probability times the total number of decisions, *d*, under evolution:

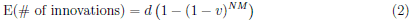

As *NM* grows, so too does the rate at which innovations arise in the population. The effect is largest if *υ* ≪ 1*/*(*NM*), so that an innovation for some decision is unlikely to arise in any given generation. In that case, *P* is approximately proportional to the population size, so

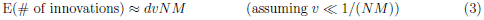

In contrast, when the population or innovation rate is large such that the inequality *υ* ≪ 1*/*(*NM*) is no longer true, then increasing *NM* has a very small effect on the total innovation rate. This is because *P* quickly approaches one; the innovation is already likely to arise *somewhere* in the population, so increasing *NM* has little effect.

### 3.2 Total population size has little effect on the spread and survival of innovations

Total population size has relatively little effect on the subsequent survival and spread of new innovations. To see this, consider the expected growth rate of a brand new innovation. Since the learning bias modeled here constitutes strong cultural selection, this initial growth rate may be the most important quantity in determining whether a new innovation will survive. Assume that other decisions are fixed for either 0 or 1, so that there is no other skill variation in the population. Let *q* = 1*/*(*NM*) be the innovation’s initial frequency in the entire population. Then the expected frequency in the next generation, *q’*, is

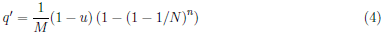

The expected growth rate upon innovation, λ, is found by dividing by *q* = 1/(*NM*), so

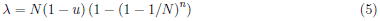

*M* drops out, showing that the number of subpopulations has no effect on the expected initial survival of new innovations. Note, though, that population density (the size of sub-populations), *N, does* affect the spread of new innovations. This effect is stronger if *n* is near *N*, i.e. if social learners can sample and accurately judge the skill of most people in the subpopulation. This partly explains why previous models (Henrich, 2004; Powell et al., 2009), which assumed *n* = *N*, found such a strong effect of interaction population size. The effect of population density is weak if *N* ≫ *n*, so that learners only learn from a small subset of a local subpopulation - say, family and close friends. In that case, λ *≈* (1 *− u*)*n*, so population density has a negligible effect on the survival of new innovations.

This derivation assumes that there is no other variation in skill in the population, so that the person with the new innovation would necessarily be the most skilled person in the subpopulation, and therefore would be copied whenever he is sampled. In general, there is likely to be skill variation present in the population for other decisions. This reduces the probability that the new innovation will be present in a highly skilled person, which ultimately reduces its probability of spreading. I doubt, however, that accounting for this process would alter the conclusion that *M* has little to no effect on the survival of new innovations.

### 3.3 Population size, equilibrium skill, and the rate of evolution

What is the overall effect of population size on the model? The above results suggest the following conclusions. First, suppose that enough time has passed for all innovations to be present in the population at some non-zero frequency. In this case, equilibrium depends on the balance between cultural “selection,” which increases mean skill, and the transmission error rate, which decreases it. Equation (5) suggests that selection depends largely on *n* and, to a lesser extent, *N*. Thus, total population size should have little affect on this equilibrium, except when *NM* is so small that drift rivals the strong effects of selection and transmission error. On the other hand, prior to reaching equilibrium, population size can strongly affect the rate of cultural evolution. If the innovation rate is very small, then large populations will improve faster because they innovate more often.

Thus, the importance of population size depends on the rate of innovation in the population and, to some extent, the timescale of interest. If innovations are rare enough to preclude populations from reaching equilibrium in historical time, then population size will correlate with skill level at any given time. If, instead, populations have reached equilibrium, then population size will have little effect. Interestingly, these two conditions resemble alternative models of genetic evolution: one in which long-term adaptation is limited by rare beneficial mutations and another in which equilibrium is determined by a mutation-selection balance. Similar conclusions apply in those models as do here.

## 4 Simulation results

### 4.1 Innovation-limited skill evolution: population size important

Figure 1a shows the rates of cultural evolution when innovation is very rare. The per-decision innovation rate is 1e-6, so *υ* ≪ *NM* in each case. Populations containing more subpopulations evolve high skill more quickly, simply because they innovate more often. Figure 1 only considers populations of total size 150 to 5,000, but the pattern holds for larger populations as long as the innovation rate is suitably small. Note, though, that the effect diminishes for larger *M*: the difference between *M* = 30 and *M* = 100 is small compared to the difference between *M* = 3 and *M* = 10.

**Figure 1:**
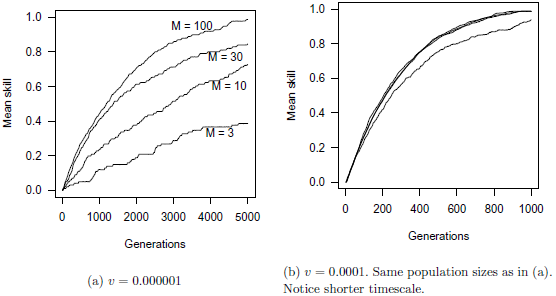
Examples of cultural evolution under varying *M*. *N* = 50, *n* = 5, *d* = 100, *m* = 0.01, *u* = 0.01. All populations eventually evolve to an equilibrium *ẑ ≈* 0.99.

Figure 1b shows the same populations when the innovation rate is 1e-4, so that the inequality *υ* ≪ *NM* no longer holds. Except for *M* = 3 (where total population size is 150), the rates of evolution are virtually indistinguishable. This is true despite the shorter timescale of figure 1b.

On the timescale of figure 1a, mean skill improvement occurs in a stepwise fashion, as can be seen by close inspection of the plot. Long periods of cultural stasis (between *∼*10 to *∼*100 generations, depending on *M*) surround short bursts of growth, during which innovations quickly spread. All populations converge on about the same equilibrium (see next section), so *M* still has little effect asymptotically. On this timescale, however, equilibrium may be so distant as to be irrelevant. If the generation time for cultural learning is equal to that of genetic reproduction (*∼*20 years), then figure 1a spans *∼*100,000 years - much longer than modern human populations have been separated. If the innovation-limited model is accurate for some skills or technologies, then it is the transient dynamics depicted in figure 1a, not the long-term equilibria, that matter.

### 4.2 Equilibrium skill level: population size not important

When innovations are not very rare, the population approaches equilibrium relatively quickly. Population size usually has negligible effect in this case. Figure 2a shows, for example, that increasing network size from five to ten has a larger effect than increasing population size from 100 to 10,000. This is a consistent finding: large changes in population size rarely alter *ẑ* by more than a few percent. Thus, small inter-group variation in sociality and communicativeness likely overshadow the effect of total population size on equilibrium skill level.

**Figure 2:**
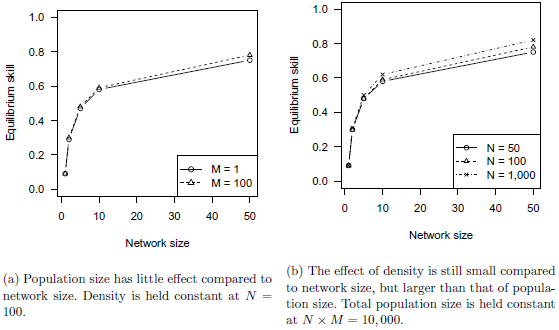
Effects of network size compared to population size and density. *d* = 100, *m* = 0.02, *v* = 0.05, *u* = 0.005.

Population density has a larger effect than population size, as predicted by equation (5). Figure 2b shows that increasing *N* from 50 to 100 has a noticeable effect on *ẑ*. This increase in density by a factor of two has about the same effect as increasing total population size by a factor of 100. As *N* grows, however, the effect becomes smaller: increasing *N* from 100 to 1,000 has about the same effect as increasing from 50 to 100. Equation (5) predicts this diminishing effect. In any case, population density is largely trumped by network size and transmission error rate.

The appendix shows the effect of population size on equilibria under various other parameter sets. They largely tell the same story: equilibrium skills levels depend much more on network size, transmission error, and innovation rate, than on either population size or density.

### 4.3 The effect of “connectedness”

Others (Henrich, 2004; Powell et al., 2009) have argued that population “connectedness,” e.g. through increased migration or trade with other populations, should have essentially the same effect as population size. A cultural group could be small, but if it frequently interacts with many other groups, then it might enjoy the benefits of a larger population size.

What affect does migration rate have here? Its primary effect is on the rate at which innovations spread between subpopulations, meaning that it is most important when innovations are rare. To see this, first imagine a metapopulation in which there is no migration at all between subpopulations. Then the rate of cultural evolution is simply equal to the mean of a single population of size *N*, since each subpopulation must innovate on its own. If innovation is rare, then subpopulations will evolve much more slowly than if they were connected by migration. If innovation is not very rare, then each subpopulation approaches equilibrium relatively quickly (figure 1b), so migration has little effect.

Simulation suggests that, as in genetic models, the effect of migration depends largely on the size of *Nm*. If *Nm* ≪ 1, then increasing migration has a large effect on rate of evolution. If *Nm* is not very small, populations behave as if effectively united, so increasing migration rate has little effect. Figure 3 shows cases where *Nm* ranges from 0 to 5. At the upper range, as *Nm >* 1, the effect of increasing *m* by an order of magnitude is small. Otherwise, the effect is large. Notice that *Nm* is simply the expected number of migrants a subpopulation exchanges with others, each generation. Thus, as long as subpopulations exchange *∼*1 migrant every generation, then changes in connectedness will have little effect. Note that this model assumes island-style migration, in which each subpopuation has equal probability of migrating to each other. A stepping-stone model is probably more realistic for humans. Such models usually show higher divergence between populations (Charlesworth and Charlesworth, 2010, ch. 7), so it is possible that migration could be important under a wider range of parameters than found here (see Powell et al., 2009).

**Figure 3:**
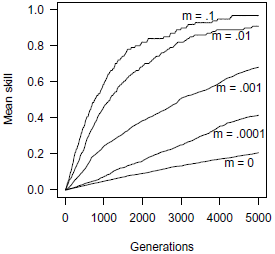
Example effects of migration on skill evolution when innovation is rare. *N* = 50, *n* = 5, *d* = 100, *u* = 0.01, *v* = 1e-6. All populations eventually evolve to an equilibrium ẑ ≈ 0.99.

## 5 Application to the current empirical debate

Although the model treated here could apply to any cumulatively evolving cultural trait, empiricists have focused mostly on technological evolution. My results suggest that the effect of population size depends on (1) whether the total innovation rate is the limiting factor on technological evolution, and (2) whether populations are near equilibrium. If technological growth proceeds slowly, by the discovery of very rare improvements (by genius or accident) which readily spread to the rest of the population, then population size may well predict technological complexity in reality. If this is a good model, then ethnographically observed levels of technology are only snapshots of a longer evolutionary process.

If, instead, innovation is not very rare, e.g. if a new innovation is likely to arise every few generations, or populations are near equilibrium, then population size should have little effect. Rather, network size, transmission error rate, and innovation rate should be most important. These parameters likely vary across different human groups. Cultures that better facilitate sociality, information sharing, and effective teaching will better be able to maintain complex technologies and skills (Henrich, 2010).

Of course, either model may be appropriate, depending on the region, the technology of interest, and the timescale. Innovations for complex or nonintuitive technologies are probably much rarer than for simpler technologies, and transmission error much more frequent. This may help to explain the ambiguous empirical support for the theory: some claim little to no effect of population size (Collard et al., 2005, 2013a; Read, 2008, 2012), while others find a strong effect (Kline and Boyd, 2010; Collard et al., 2013b). Collard et al. (2013a) surmises that population size has the *potential* to affect technological complexity, but probably will not *always* do so. My results here offer some mechanistic insight as to why the effect of population size would be so heterogeneous.

I suspect, however, that the discrepant findings do not reflect variation in actual technological processes, but rather reflect imperfect, divergent methods. For example, no study measures population density, village/band size, or anything akin to social network size. These have distinct affects from population size, but they are probably all positively correlated: large populations will, other things being equal, tend to be dense, and people in denser populations probably have more social contacts. It is unclear, then, whether an observed relationship between population size and technological complexity implies an actual effect of population size *per se*. Furthermore, different studies consider very different regions, often without accounting for population history or inter-group contact. Studies that limit attention to specific island regions (Kline and Boyd, 2010; Read, 2012) probably have better control over these variables, but they also preclude generalization to other regions and larger geographical areas. Finally, technological complexity itself also affects population growth, implying a reverse causation that none of our models or statistical studies have considered.

Thus, the debate remains whether population size has an important effect on technological evolution. For future empirical studies, researchers should use measures of population density and network size, so as to better distinguish their effects from total population size. Network size can, perhaps, be estimated by village or band size, at least when these are small.

## A The effect of ***M*** on equilibrium skill for various parameters

I found equilibrium skill levels, *ẑ*, by fitting a first-order autocorrelated normal model on each time-series, after convergence to a stationary distribution. I round estimates to the nearest 0.01; these are accurate in each case with at least 95% confidence (and usually much more). Tables show first-order interactions of *M* with each other variable. For each table, the non-focal parameters are kept at baseline values. These are *N* = 100, *M* = 50, *n* = 10, *m* = 0.01, *u* = 0.05, *v* = 0.001, and *d* = 100.

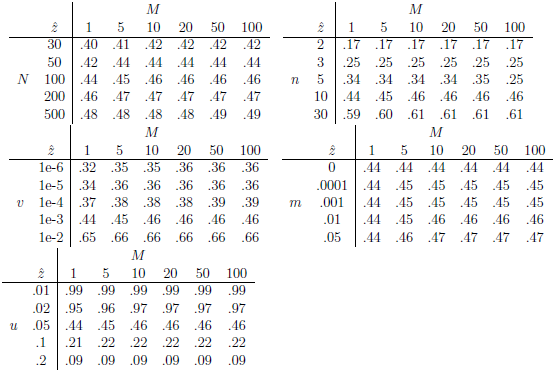

